# Plasmacytoid dendritic cells mediate CpG-ODN induced increase in survival in a mouse model of lymphangioleiomyomatosis

**DOI:** 10.1101/2023.02.06.527331

**Authors:** Mayowa M Amosu, Ashleigh M Jankowski, Jacob C McCright, Bennett E Yang, Juan Grano de Oro Fernandez, Kaitlyn A Moore, Havish S Gadde, Mehul Donthi, Michele L Kaluzienski, Vedanth Sriram, Katharina Maisel

## Abstract

Lymphangioleiomyomatosis (LAM) is a devastating disease primarily found in women of reproductive age that leads to cystic destruction of the lungs. Recent work has shown that LAM causes immunosuppression and that checkpoint inhibitors can be used as LAM treatment. Toll-like receptor (TLR) agonists can also re-activate immunity and the TLR9 agonist, CpG-ODN, has been effective in treating lung cancer in animal models. Here we investigate the use of TLR9 agonist CpG-ODN as LAM immunotherapy in combination with checkpoint inhibitor, anti-PD1, standard of care rapamycin and determine the immune mechanisms underlying therapeutic efficacy. We used survival studies, flow cytometry, ELISA, and histology to assess immune response and survival after intranasal treatment with CpG-ODN in combination with rapamycin or anti-PD1 therapy in a mouse model of metastatic LAM. We found that local administration of CpG-ODN enhances survival in a mouse model of LAM. We found that a lower dose led to longer survival likely due to fewer local side effects but increased LAM nodule count and size compared to the higher dose. CpG-ODN treatment also reduced regulatory T cells and increased the number of Th17 helper T cells as well as cytotoxic T cells. These effects appear to be mediated in part by plasmacytoid dendritic cells (pDCs), as depletion of pDCs reduces survival and abrogates Th17 T cell response. Finally, we found that CpG-ODN treatment is effective in early stage and progressive disease and is additive with anti-PD1 therapy and rapamycin. In summary, we have found that TLR9 agonist CpG-ODN can be used as LAM immunotherapy and effectively synergizes with rapamycin and anti-PD1 therapy in LAM.

## INTRODUCTION

Lymphangioleiomyomatosis (LAM) is a progressive rare lung disease, primarily affecting women who are diagnosed in their 20-40s (1). LAM patients can experience a range of debilitating symptoms including shortness of breath, lung collapse, and eventual respiratory failure. LAM is characterized by the persistent abnormal growth of smooth muscle-like cells (LAM cells) and the formation of thin-walled cysts predominantly in the lungs, with involvement of the lymphatic system and kidneys. LAM is caused by inactivating mutations in tumor suppressor genes tuberous sclerosis complex 1 and 2 (TSC1 / TSC2) that negatively regulate cell growth through inhibition of a signaling protein complex called mechanistic target of rapamycin (mTOR). Currently, sirolimus (rapamycin) is the only FDA approved drug for LAM (2), which inhibits the constitutively activated mTOR complex thus slowing disease progression (3,4). However, rapamycin does not promote disease regression or lung regeneration (5,6). Furthermore, patient responses to rapamycin are incomplete and not always well tolerated for life-long use (5). Therefore, there is an urgent need for curative treatments that can eliminate LAM nodules and restore healthy lung function in patients suffering from LAM.

LAM is designated as a neoplastic growth, and recent research suggests it has similarities to cancer (7). Immunosuppression is a common feature of many cancers, including lung cancers (8,9), which over time conditions immune cells to ignore the tumor cells thus promoting cancer progression (10–12). In human LAM tissues, a lack of T-cell infiltration suggests immunosuppression due to unresponsiveness of lymphocytes to LAM antigens (13). LAM lungs also typically have elevated levels of secreted factors like monocyte chemoattractant protein 1 (MCP-1) (14), which may contribute to recruitment of immunosuppressive cells (15,16). Prior work by us and others has also shown that human LAM nodules have elevated expression of PD-L1, an immune checkpoint inhibitor that causes T cell suppression, and LAM causes overexpression of checkpoint molecules PD-1 and CTLA-4 on T cells in a mouse model (17,18). Anti-PD1/PD-L1 and anti-CTLA-4 therapies were shown to enhance survival in mouse models of LAM (17,18), suggesting that countering immunosuppression in LAM could be an effective strategy for treating LAM.

In addition to the previously investigated checkpoint inhibitor treatments, reducing immunosuppression in LAM could be accomplished by reactivating antigen presenting cells (APCs) in the local tumor environment. Recent studies investigating targeted activation of APC’s in cancer show enhanced anti-tumor immunity through improved T cell responses (19), and increased responsiveness of immunosuppressed myeloid cells to immunotherapy (20). Local tumor treatments inducing APC activation have also been shown to stimulate systemic anti-tumor immunity and regression of distant tumors (21). Treatments that mimic microbial pathogens to stimulate toll-like receptors (TLRs) on APCs have been explored for this purpose. When engaged through TLR activation, APCs like dendritic cells and macrophages secrete inflammatory cytokines and promote the differentiation of cytotoxic T cells, reduce regulatory T cells, and can lead to increases in pro-inflammatory helper T cells (22,23). Several TLR agonists have received FDA approval for cancer treatment. For example, BCG is a TLR4 agonist that is used as immunotherapy for bladder cancer and there are at least 8 additional clinical trials are investigating TLR adjuvants as cancer therapeutics (NCT04364230, NCT04819373, NCT04278144, NCT03486301, NCT04840394, NCT02668770, NCT05607953, NCT04935229). TLR9 agonists, like CpG oligonucleotides (CpG-ODN), have been explored as therapy in multiple cancers including non-small cell lung cancer and are FDA approved vaccine components (24–26). TLR9 agonists have also been shown synergize with immune checkpoint inhibition, leading to increased therapeutic responses (27–31). Given the current efforts to leverage the TLR9 agonist CpG-ODN in cancer therapy, we investigated its potential for therapy in LAM. We assessed the therapeutic efficacy of CpG-ODN and the immune mechanisms involved in its therapeutic action, and determined efficacy in combination with checkpoint inhibition or the standard of care rapamycin treatment.

## METHODS

### Mice

7 to 9-week old female C57BL6/J mice (Jackson Laboratory) were used for all studies. Study protocols were approved by the UMD Institutional Animal Care and Use Committee (IACUC).

### LAM induction

TSC2-null kidney-derived epithelial tumor (TTJ) cells, were provided by the Krymskaya Lab at the University of Pennsylvania (17). TTJ cells (5.0 x 10^5^) were administered in 100 µL of PBS via tail vein injection to induce LAM disease in 8-9 week old female mice. Lung lesions typically develop within 3-7 days.

### Survival Studies

Mice received CpG-ODN class B (5 μg/10 μg) 2x/week (Invivogen, tlrl-1826-5), in 50 µL of PBS, intranasally starting 4 days post LAM induction. Mice receiving combination therapy, received intranasal CpG-ODN and 300 µg anti-PD-1 antibody (BioXCell, BE0146-A0) or 1 µg/g Rapamycin (Selleck Chemicals, S1039) in 200 µL of PBS via intraperitoneal (IP) injection 2x/week. Control groups received 50 µL of PBS intranasally or 200 µL PBS with or without 300 µg of rat IgG isotype control (BioXCell, BE0089) by IP injection. Mice were monitored for symptoms of disease and body weight changes and removed from studies based on experimental timing or humane endpoints. At endpoint, mice were euthanized and lung tissues collected for histological analysis.

### Histology

Tissues were fixed overnight in 4% PFA at 4⁰C, embedded in paraffin wax, sectioned at 5 µm, and stained with hematoxylin and eosin (H&E). LAM nodule counts, average nodule diameters, and general tissue inflammation were quantified using QuPath Imaging software.

### Flow Cytometry

After LAM induction, mice received a single intranasal treatment dose of CpG-ODN (5 μg or 10 μg) in 50 µL PBS or 50 µL of PBS as a vehicle control. At 1 and 3 days post treatment, lung tissues were perfused and collected for flow cytometric analysis. Left lung lobes were dissociated by enzymatic digestion with 1 mg/mL collagenase D (Roche, 11088882001), and 1 mg/mL collagenase IV (Worthington Biochemical, LS004188) in DMEM with 5% FBS for 1 hour. Single cell suspensions were stained to identify eosinophils (CD64-, Siglec F+, Ly6G-), neutrophils (CD64-, Siglec F-, Ly6G+) dendritic cells (CD11c+, MHC-II^high^), plasmacytoid dendritic cells (CD11c+, PDCA-1+, MHC-II^high^), alveolar macrophages (CD64+, SiglecF^high^, CD11b-), interstitial macrophages (CD64+, SiglecF^low^, CD11b+, MHC-II+), B Cells (CD19+ or B220+), T Cells (CD3+, CD4+ or CD8+), regulatory T Cells (CD3+, CD4+, FoxP3+), Th1 T cells (CD3+, CD4+, FoxP3-, Tbet+), Th2 T cells (CD3+, CD4+, FoxP3-, Gata3^hi^), and Th17 T Cells (CD3+, CD4+, FoxP3-, ROR-γt+). Cells were analyzed by flow cytometry using a BD FACSCelesta (BD Biosciences) and FlowJo Version 10 (FlowJo LLC). Gating strategies can be found in the data supplement (**Fig E1**).

### ELISA

For cytokine analysis, lungs (smallest right lobe) were snap-frozen and stored at −80°C. For tissue homogenates, snap-frozen lungs were homogenized at 4°C at 20% w/v in protein extraction reagent (T-PER, Thermo Fisher Scientific) containing Halt™ Protease Inhibitor Cocktail (Thermo Fisher Scientific). Lung tissue homogenates were centrifuged at 15,000 g for 10 min at 4°C. Supernatants were aliquoted and stored at −80°C. Supernatants were diluted 1:10 with 1% BSA ELISA assay diluent and ELISA was performed according to manufacturer’s instruction (ELISA MAX™ Standard Sets, BioLegend).

### Statistical Analysis

Data was analyzed using Graphpad Prism (GraphPad, La Jolla, CA). Survival studies were assessed using the log-rank (Mantel-Cox) test. Unpaired Welch t tests were used to compare pairs of samples or groups. Mann–Whitney test was used to compare the groups of data that failed to satisfy parametric assumptions. Brown-Forsythe and Welch One-way ANOVA were used for comparisons between ≥3 groups. Differences were considered statistically significant when p < 0.05. *p<0.05, **p<0.01, ***p<0.001, ****p<0.0001.

## RESULTS

### Treatment with intranasal CpG-ODN increases survival in LAM

To determine the therapeutic benefit of TLR9-agonist CpG-ODN in LAM disease, we performed survival studies comparing the effect of two doses of intranasal using our metastatic mouse model of LAM. We found that intranasal CpG-ODN significantly increases survival in mouse LAM, with median survival increasing 1.4-fold from 32.5 to 45 days (**Fig 1A**). We found that a lower dose of CpG-ODN (5mg) resulted in higher survival than higher dose CpG-ODN (10mg), with median survival increasing further from 45 to 60 days, an almost 2-fold increase compared to untreated mice. We conducted a cross sectional study after 3 weeks (Day 22) to assess treatment effect on LAM nodule formation. Histological analysis showed that treatment with CpG-ODN reduced LAM nodules in both size and number in treated vs untreated lungs (**Fig 1B**). Higher dose CpG-ODN (10ug) significantly decreased the nodule count, more than the lower dose (5ug), which was surprising given the improvement in overall survival with lower dose CpG-ODN (**Fig 1C**). However, the overall abnormal tissue fraction is similar for both doses of CpG-ODN, even though the higher dose has a lower nodule burden (**Fig 1C**), suggesting that negative side effects of CpG-ODN may contribute to the increased morbidity with higher dose CpG-ODN. In addition, we found that treatment with higher dose CpG-ODN results in similar levels of immune cell infiltration to untreated LAM, while treatment with lower dose CpG-ODN results in less CD45+ cells than untreated LAM (**Fig E2**).

**Figure 1.**
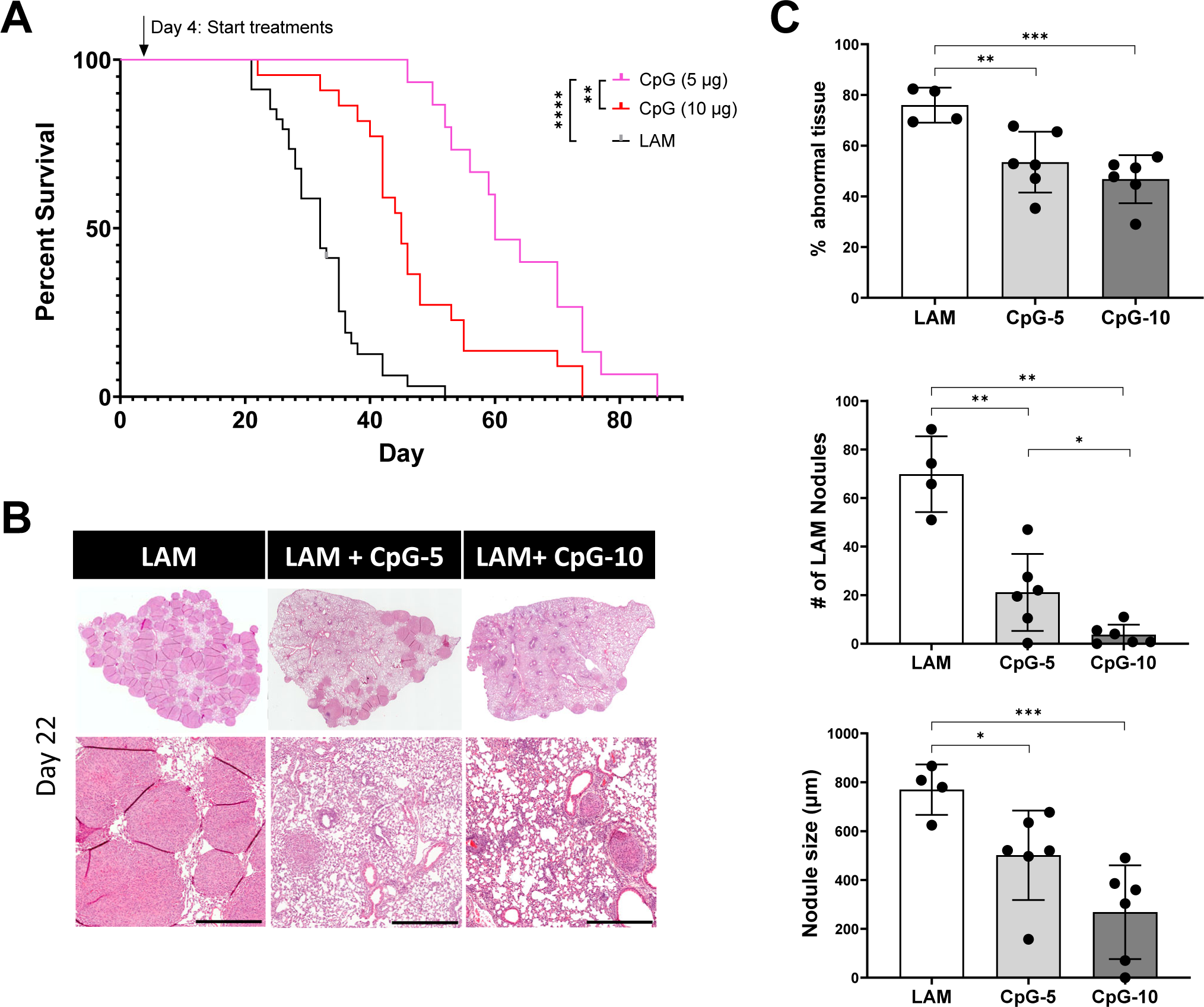
Intranasal CpG increases survival in murine model of metastatic LAM, decreases tissue-wide inflammation and overall LAM nodule burden. **A)** Aggregate data from three survival experiments LAM n=34, CpG (10ug) n=22, CpG (5ug) n=15. **B)** Representative H&E stained histological images of mouse lungs from treatment groups: untreated LAM control, CpG 5 µg, and CpG 10 µg. Scale bar = 500 µm. **C)** Quantification of LAM nodules and Inflamed tissue regions between treatment groups 22 days post LAM induction and after 6 total treatment administrations. 3 sections analyzed per mouse. Statistical analysis was performed using log-rank test (survival studies) and one-way ANOVA (histological analysis). * p<0.05, ** p<0.01, *** p<0.001, **** p<0.0001.

### Intranasal CpG-ODN modulates Immune cell infiltration in LAM Lungs and shifts T Cells toward inflammatory phenotypes and reduces regulatory T cells

We next sought to determine the immune mediators of CpG-ODN treatment that could be responsible for the treatment efficacy. We assessed immune cell subsets early during intervention at day 5 and later at day 22. On day 5, populations of eosinophils, alveolar macrophages, dendritic cells, and B and T cells were significantly decreased at both low and higher dose CpG-ODN. In contrast, neutrophils and interstitial macrophages, and monocytes were increased in response to CpG-ODN treatment (**Fig 2A, Figure E3**). By day 22, CpG-ODN treatment in LAM mice resulted in decreased cell counts in all identified immune populations, except B cells and interstitial macrophages, when compared to counts in untreated LAM controls (**Fig 2B**). Between Day 5 and Day 22, untreated mice had the greatest expansion of interstitial macrophages and cDCs, while CpG-ODN treated mice had greatest expansion in cDCs and T Cells (**Fig E3-4**). We also found up to 5-fold increases in T and B cell numbers in CpG-ODN treated mice compared to up to 10 fold increases in disease controls (**Fig E3-4**). B cells in untreated mice were the only cell population with little expansion from Day 5 to 22. In CpG-ODN treated lungs, there was no expansion in eosinophil, alveolar macrophage and monocyte populations (**Figure E3**). The 10-fold expansion of CD8+ T cells and 7.5-fold increase in CD4+ T cells **(Fig E4)**, was correlated with an increase in T cell chemoattractant CCL21 in CpG-ODN treated mice **(Fig E5).** This suggests that CpG-ODN stimulates chemokine release in the tissue and correlates with increased lymphocytes in the lungs.

**Figure 2.**
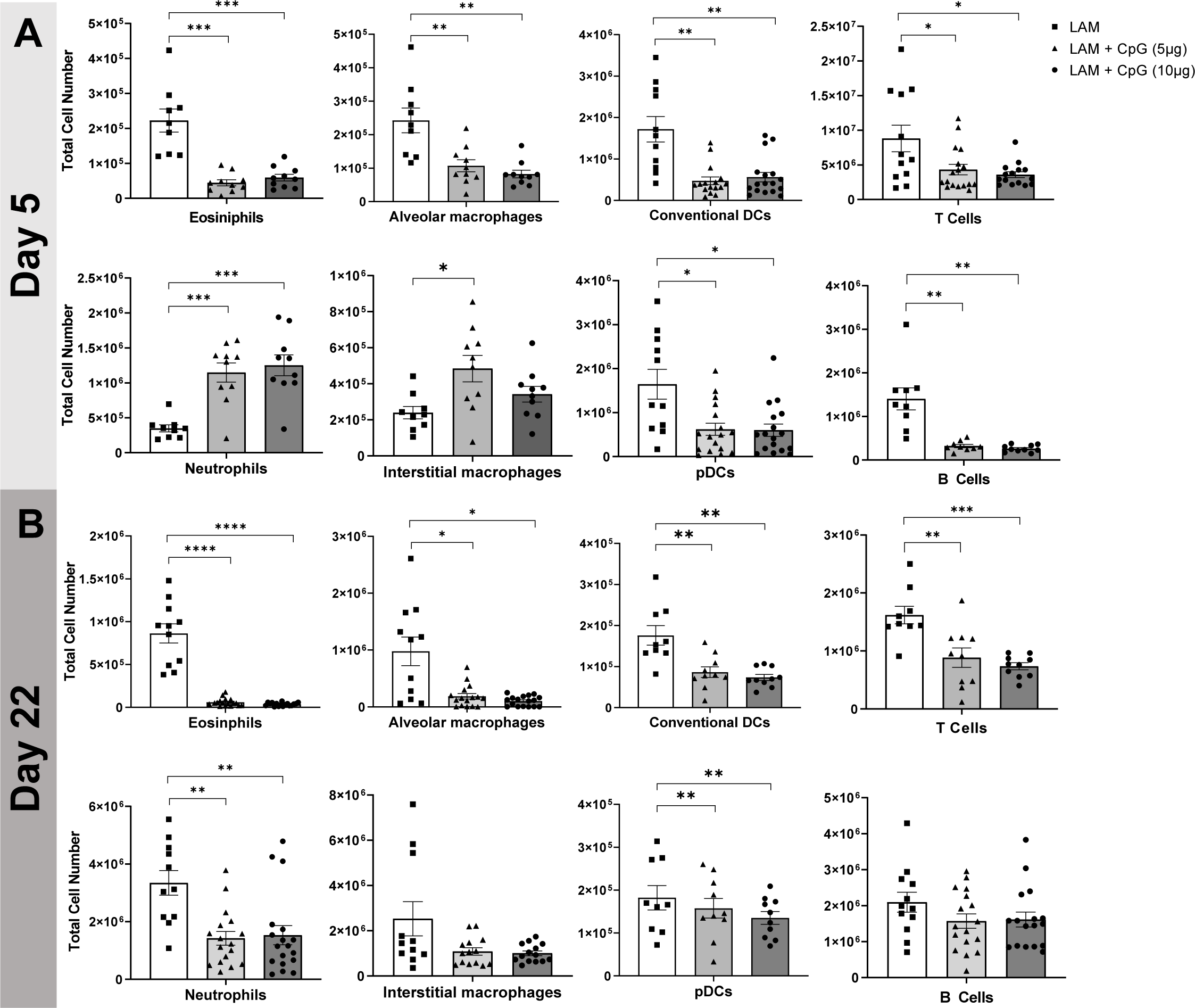
CpG treatment modulates immune cell infiltration in LAM. Data shows total cell numbers in the left lung lobe of eosinophils, neutrophils, alveolar macrophages, interstitial macrophages, conventional DCs, plasmacytoid DCs, T cells and B cells at **A)** day 5 or **B)** day 22 after LAM induction and 1 or 18 days respectively after CpG treatment has started. Data is representative of n= 2+ experiments and n>8 mice. Statistical analysis was performed using one-way ANOVA. * p<0.05, ** p<0.01, *** p<0.001, **** p<0.0001.

While we saw an overall reduction in CD4+ and CD8+ T cell numbers with CpG-ODN treatment in LAM lungs (**Fig E6**), we found a shift in the activation and skewing of both CD8+ and CD4+ T cells in the lungs. Flow cytometry analysis revealed that CD8+ T cells have increased expression of activation marker ICOS (**Fig 3A**) and the proportion of effector memory CD8+ T cells is significantly increased with CpG-ODN treatment (**Fig 3B**), while naïve CD8+ T cells are reduced with the highest dose CpG-ODN. This suggests that CpG-ODN treatment activates cytotoxic CD8+ T cells, which was further confirmed by the increase in CD8+ T cells producing cytotoxic molecules and cytokines including granzyme B, IFN-γ, TNFα, and perforin (**Fig 3C**). We also found signs of reduced immunosuppression as indicated by the significantly decreased regulatory T cell numbers and reduced ICOS+ regulatory T cells (**Fig E6B**) in CpG-ODN treated mice (**Fig 3D**). Furthermore, we found increased proportions of Th1 and Th17 cells and increased ICOS+ Th17 cells (**Fig 3D**), suggesting that T cells in CpG-ODN treated LAM lungs are polarized towards more proinflammatory phenotypes. Cytotoxic T cells and Th1 and Th17 cells are known to produce pro-inflammatory IFN-γ, and we found increased levels of IFN-γ in LAM lung tissues after CpG-ODN treatment (**Fig 3E**), suggesting increased cell-mediated cytotoxicity. We also found an increase in IL12/23 in LAM lung tissues after CpG-ODN treatment (**Fig 3E**), which are important mediators of CD4+ T cell differentiation, with IL-23 pushing Th17 differentiation, which is consistent with the increased proportion of Th17 cells we also found in LAM lungs after CpG-ODN treatment.

**Figure 3.**
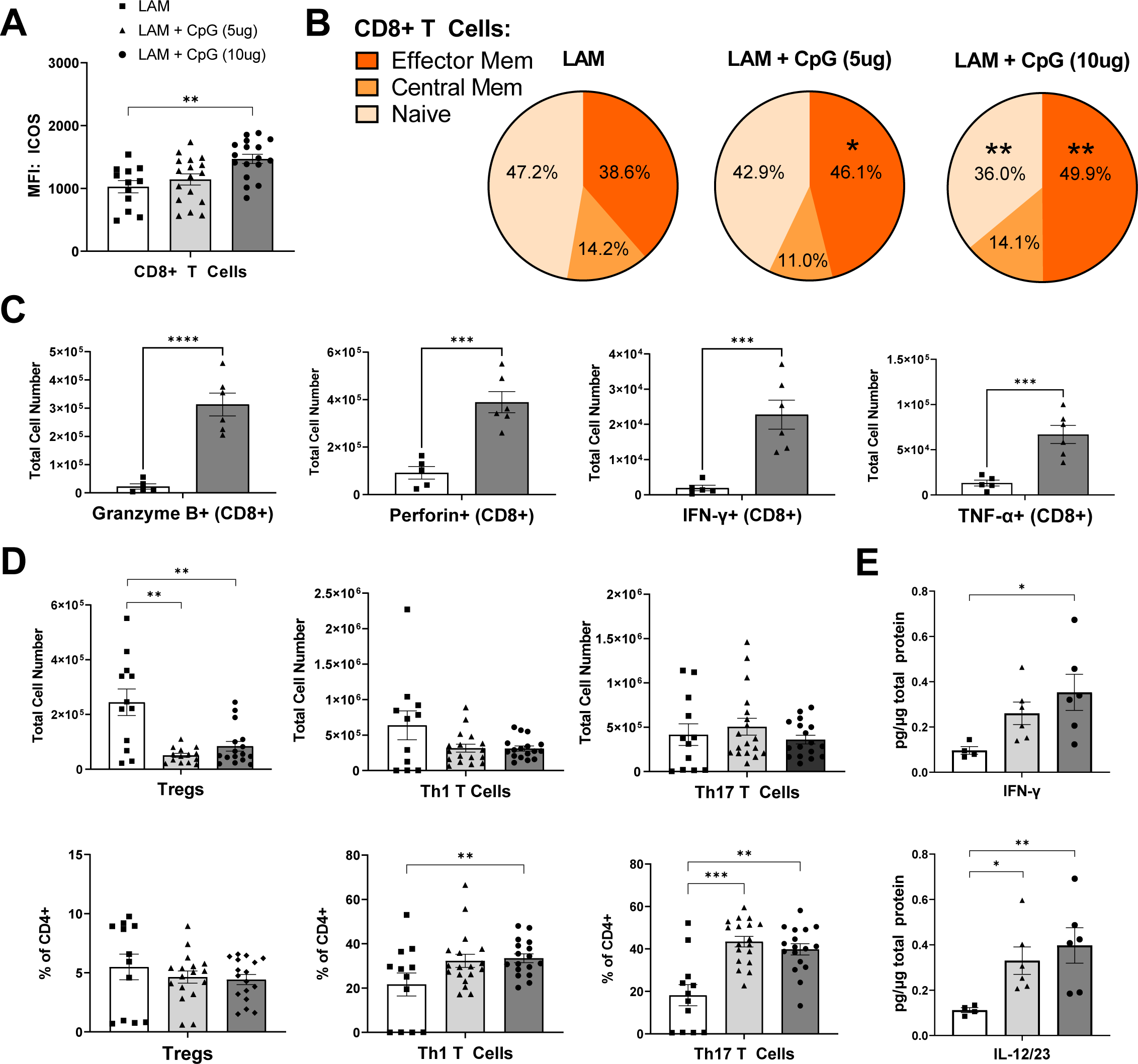
Helper T cells in CpG treated LAM lungs shift away from immunosuppressive phenotypes and CD8 T cells have increased effector functions. Data shows T cell populations in the left lung lobe 22 days after LAM induction. **A)** MFI of ICOS expression in CD8+T Cells. **B)** Effector memory, central memory, and naïve CD8+ T cell subsets. **C)** Intracellular cytokines stained in CD8+ T cells. **D)** CD4 helper T cell subsets: total cell numbers and % of CD4+ cells. **D)** IFNy and IL12/23 cytokine levels in LAM lungs. Data is representative of n= 2+ experiments and n>5 mice. Statistical analysis was performed using one-way ANOVA. * p<0.05, ** p<0.01, *** p<0.001, **** p<0.0001.

### Therapeutic efficacy of CpG-ODN is mediated by plasmacytoid dendritic cells

Plasmacytoid dendritic cells (pDCs) express high levels of TLR9 in both mouse and human lungs (32–36). Therefore, we hypothesized that pDCs would mediate CpG-ODN treatment efficacy by activating and skewing T cells. We depleted pDCs using anti-PDCA-1 antibodies (**Fig E7**) and examined treatment outcome and immunity. We found that pDC depletion in CpG-ODN treated mice with LAM significantly reduced survival, with a decreased median survival from 60 days to 56 days (**Fig 4A**). The effect of CpG-ODN treatment on survival remains significant despite depletion of pDCs, suggesting that although pDCs play a part in the response that makes CpG-ODN treatment beneficial, other cell types and pDC-independent mechanisms likely contribute to this treatment effect. To assess the effect of pDCs on T cell immunity in LAM, we also assessed CD4+ T cell subsets after pDC depletion and found that the number of Treg cells remained reduced but the proportion of Th17 cells was no longer increased compared to CpG-ODN treatment alone (**Fig 4B**). Additionally, we found that pDC depletion abrogated the increase in IL-12/23 and IL-6 induced by CpG-ODN treatment (**Fig 4C**); both key cytokines that promote Th17 differentiation, suggesting that pDCs are at least in part required for inducing Th17 responses after CpG-ODN treatment in LAM. In CD8+ T cells, pDC depletion resulted in a significant decrease in the secretion of inflammatory cytokines, granzyme B, perforin, IFN-γ, and TNF-α (**Fig 4D**), suggesting that pDCs play a role in activating these cytotoxic responses stimulated by CpG-ODN-treatment in LAM.

**Figure 4.**
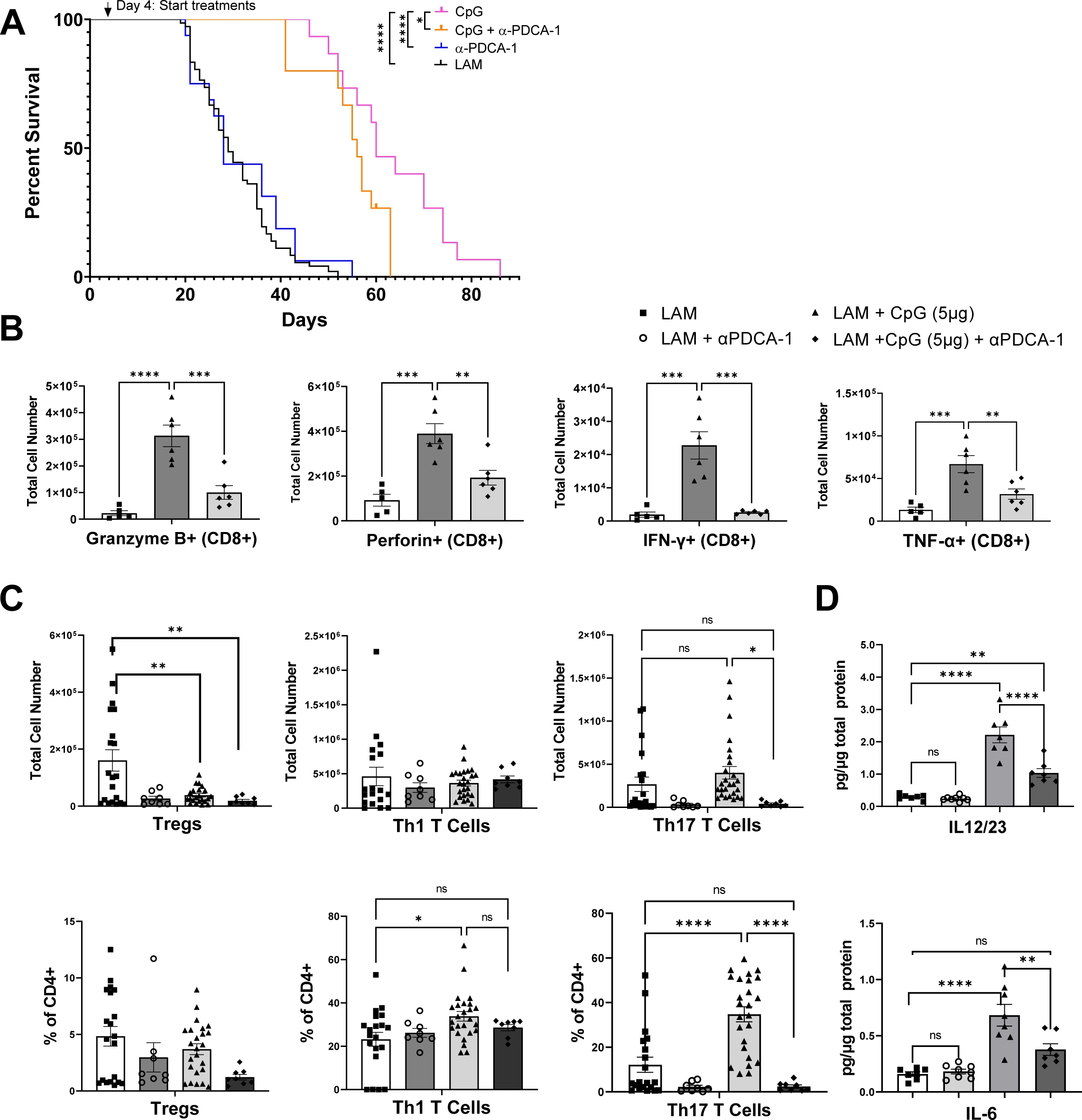
pDC depletion in CpG-treated LAM decreases survival and affects T cell responses. **A)** Survival studies showing pDC depletion reduces survival after CpG treatment. **B)** Number of CD8+ T cells producing Granzyme B, Perforin, IFN-γ, or TNF-α. **C)** CD4 helper T cell subsets including Tregs, Th1s, and Th17s: total cell numbers and % of CD4+ cells. **D)** IL12/23 and IL-6 cytokine levels in LAM lungs. Data is representative of n=2+ experiments and n>6 mice. Statistical analysis was performed using log-rank test (survival studies) and one-way ANOVA. * p<0.05, ** p<0.01, *** p<0.001, **** p<0.0001.

### CpG-ODN treatment is less effective in late stage disease

Given the encouraging results we found with CpG-ODN treatment in early LAM, and since LAM can be diagnosed at a variety of disease stages, we next evaluated CpG-ODN’s therapeutic potential in more advanced disease. We induced LAM in mice and allowed disease to develop for 14 days, before beginning intranasal treatment with CpG-ODN. We found that the previously beneficial lower dose CpG-ODN (5 µg) had no significant effect on survival (**Fig 5A**), but a high dose of CpG-ODN (20 µg) showed some therapeutic benefit (**Fig 5A**), increasing median survival from 32 days to 40 days. Despite the increased survival, we found that CpG-ODN treatment did not improve abnormal tissue fractions, nodule count, or nodule size in more advanced disease (**Fig 5B-C**). However, we did observe that nodules in CpG-ODN-treated mice showed histological signs of necrosis in LAM nodules, unlike untreated LAM controls (**Fig 5B**). We then determined the effect of CpG-ODN on the immune response in late stage LAM mice 1 and 3 days following the first intranasal treatment of CpG-ODN (days 15 and 17 after disease induction). Unlike the striking responses seen during early intervention, CpG-ODN given later in disease did not significantly affect immune cell infiltration in LAM lungs. There were no differences in total CD45+ immune cell populations between CpG-ODN treated and untreated LAM lungs (**Fig E8**). We found only a decrease in eosinophils and cDCs at day 15; all other populations of identified immune cells remained unchanged (**Fig 6**). Overall, our data suggest that CpG-ODN treatment, while increasing survival, has a minor beneficial effect in late stage disease.

**Figure 5.**
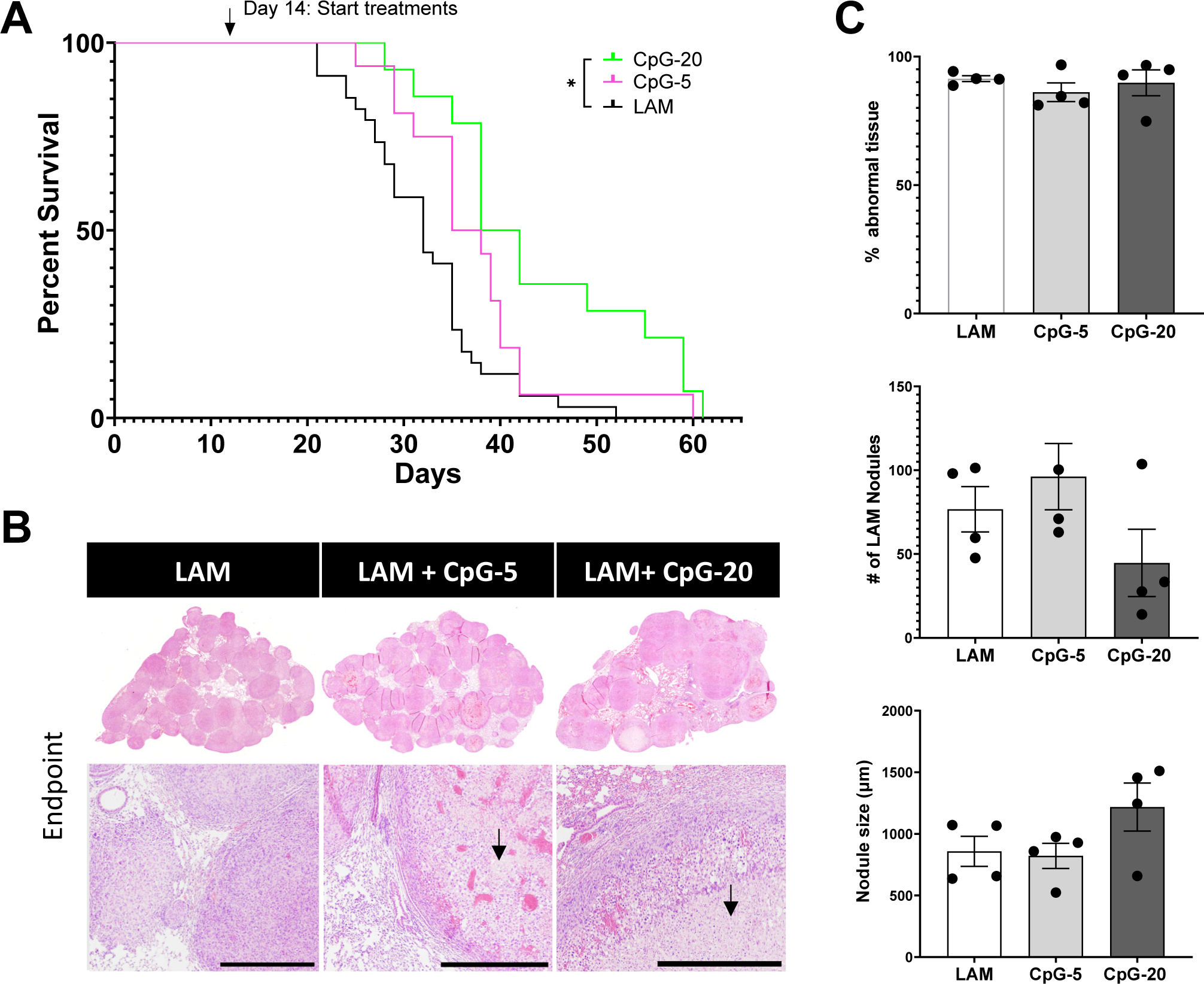
Late intervention intranasal CpG increases survival in murine model of metastatic LAM at higher dose. **A)** Survival studies showing 20 µg and 5 µg CpG treatments in late stage disease. **B)** Representative H&E images of mouse lungs from treatment groups: untreated LAM control, CpG 5 µg, and CpG 20 µg. Arrows indicate areas of necrosis. Scale bar = 500 µm. **C)** Quantification of LAM nodules and abnormal tissue regions for treatment groups of mice at symptomatic endpoints from survival studies. Data is representative of n=2+ experiments and n>4 mice. Statistical analysis was performed using log-rank test (survival studies) and one-way ANOVA (histological analysis). * p<0.05.

**Figure 6.**
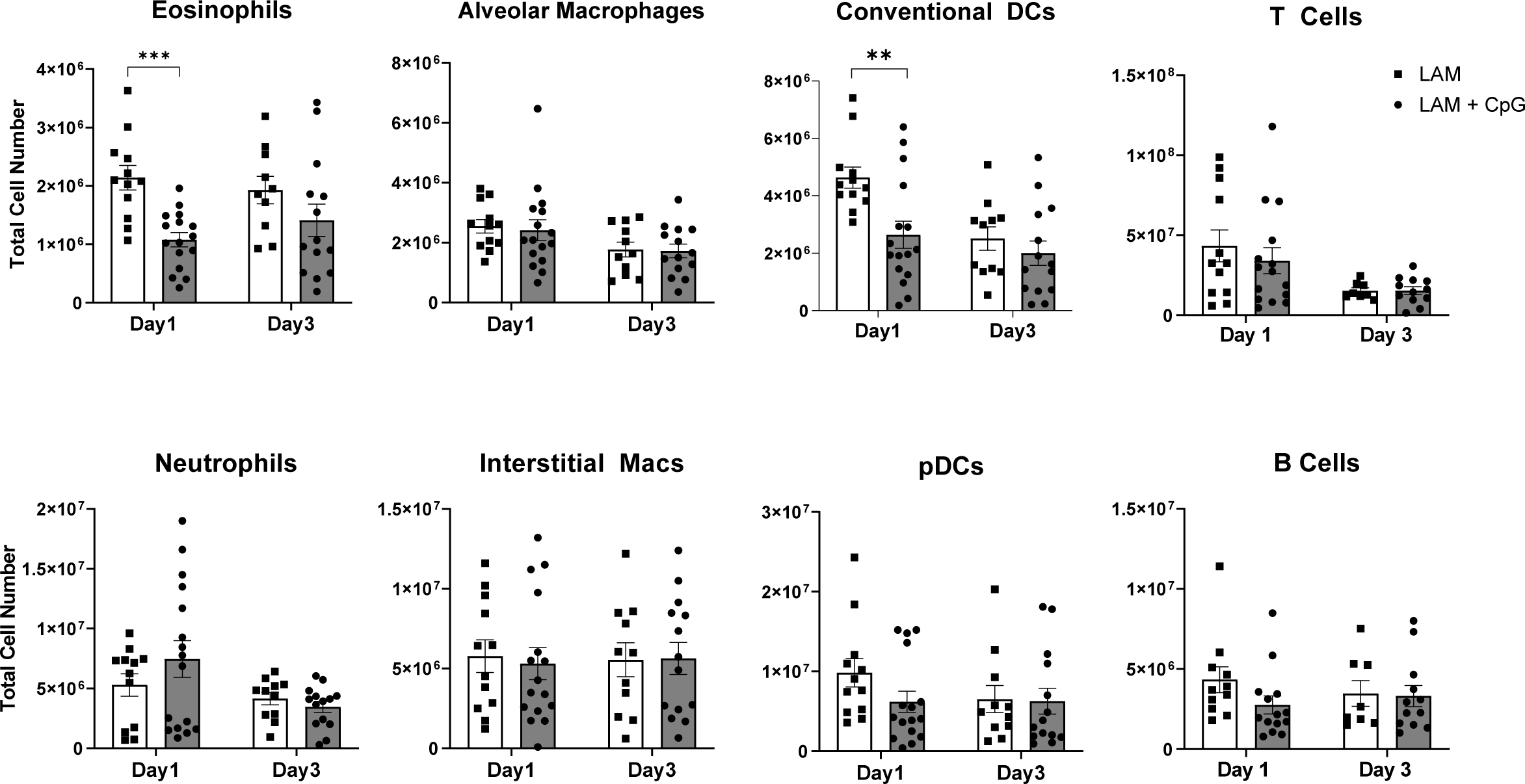
CpG treatment modulates immune cell populations in response to late intervention (starting at day 14) intranasal CpG in LAM lungs (Day 1 & Day 3 after CpG treatment). Total cell numbers of granulocytes and antigen presenting cells infiltrating the lungs 1 and 3 days after CpG treatment obtained via flow cytometry. Data is representative of n= 3 experiments and n = 8+ mice. Statistical analysis was performed using one-way ANOVA. * p<0.05, ** p<0.01, *** p<0.001.

### Combination therapy of CpG-ODN with anti-PD1 or rapamycin synergizes to increase survival in mouse LAM

Rapamycin is the only treatment currently available to LAM patients, so it is crucial to evaluate combination therapy of CpG-ODN with low-dose rapamycin for clinical translation. We found that combination CpG-ODN with low-dose Rapamycin therapy significantly increased survival in mice (**Fig 7A**) compared to CpG-ODN or Rapamycin treatments alone, increasing median survival more than 3-fold, from 29 days in untreated mice to 100 days in mice receiving combination therapy. The median survival in mice receiving monotherapies led to a 2-fold increase from 29 days to 60 days for both CpG-ODN and Rapamycin treatment groups. Histological analysis of tissues collected at endpoints from survival studies showed that combination of CpG-ODN with Rapamycin therapy significantly reduced LAM nodule counts and size, and abnormal lung tissue compared to CpG-ODN monotherapy (**Fig. 7B-C**).

**Figure 7.**
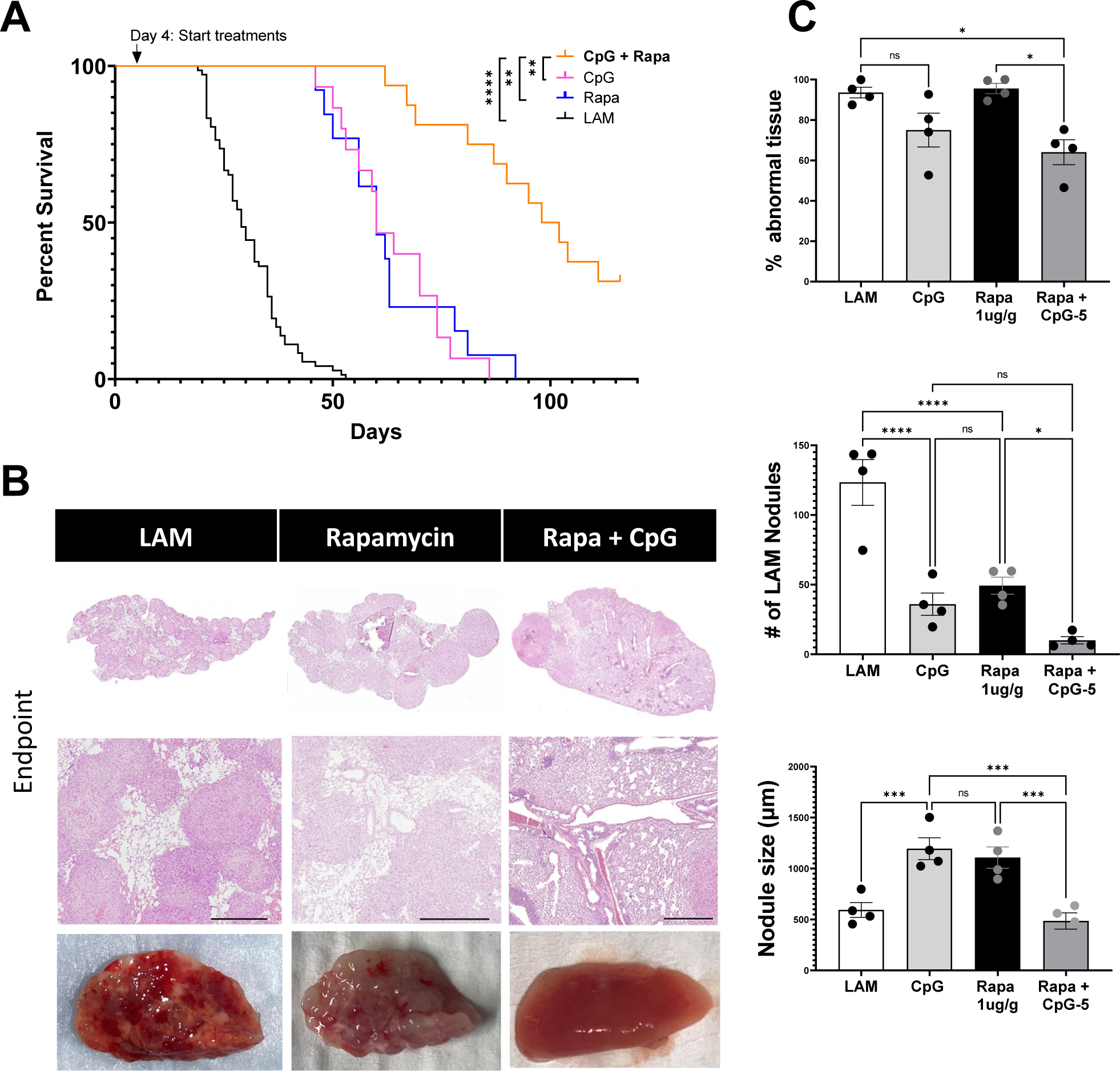
CpG treatment is synergistic with Rapamycin Therapy. **A)** Aggregate data from four survival experiments. LAM n=72, CpG (5 µg) n=15,Rapa n=13, CpG (5 µg) + Rapa n=16. **B)** Representative H&E stained histological images of mouse lungs from treatment groups: untreated LAM control, CpG, Rapa, CpG + Rapa. Scale bar = 500 µm. **C)** Quantification of LAM nodules and Inflamed tissue regions between treatment groups of mice at symptomatic endpoints from survival studies in (A). 3 sections analyzed per mouse. Statistical analysis was performed using log-rank test (survival studies) and one-way ANOVA (histological analysis). * p<0.05, ** p<0.01, *** p<0.001, **** p<0.0001.

We have also previously shown that treatment with anti-PD-1 antibody prolongs survival in a mouse model of LAM (17). CpG-ODN has been shown to synergize with checkpoint inhibition in pre-clinical cancer models (37–39) and thus we sought to evaluate combination therapy of CpG-ODN with checkpoint inhibition to determine synergistic effects in LAM. We found that combination CpG-ODN and anti-PD-1 therapy increased survival in mice (**Fig 8A**) compared to CpG-ODN or anti-PD1 treatment alone, increasing survival to 65 days for combination therapy compared to 45 and 49 days for the monotherapies, respectively.

**Figure 8.**
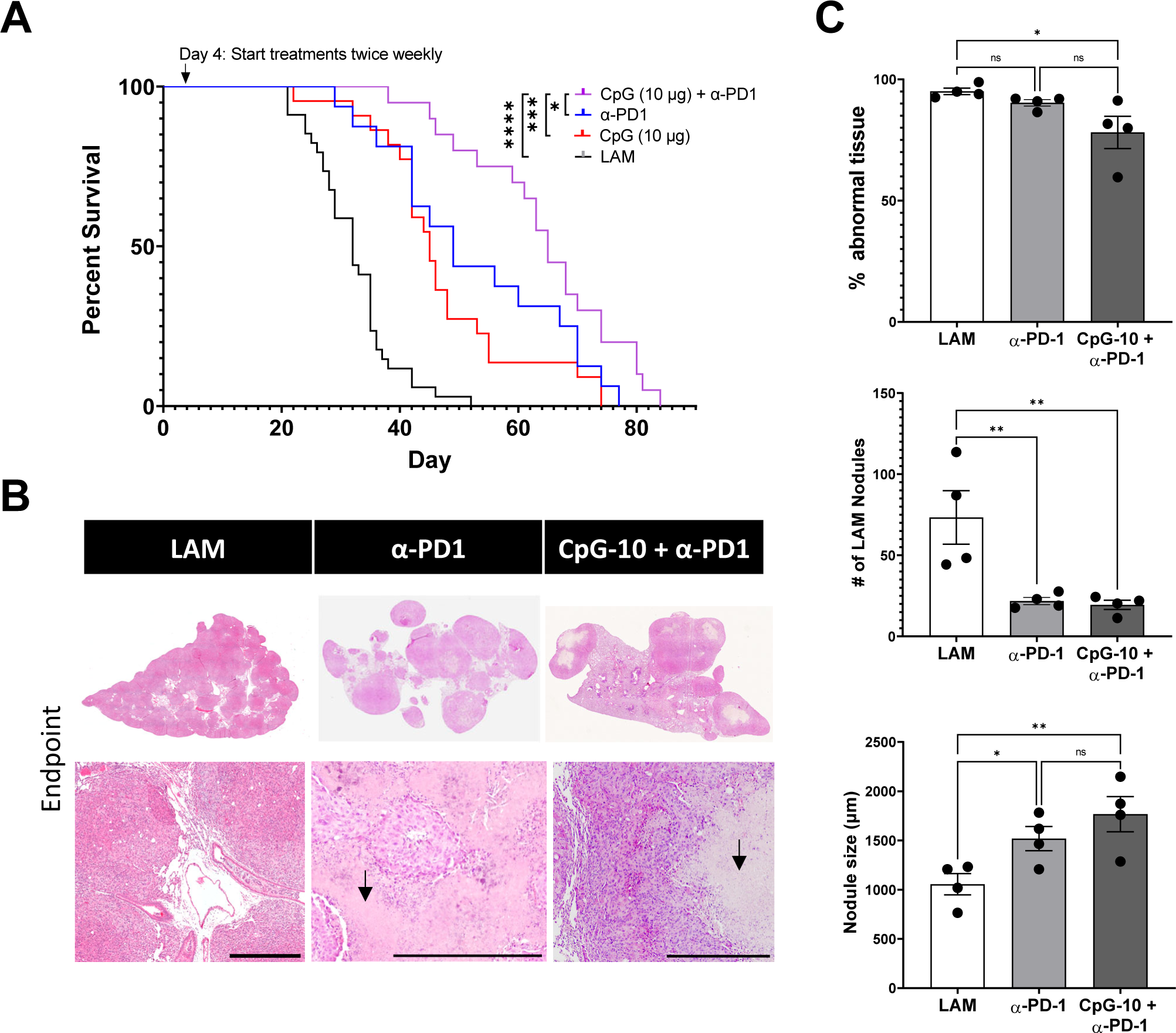
CpG treatment is synergistic with ICI Therapy. **A)** Aggregate data from four survival experiments LAM n=34, CpG (10 µg) n=22, α-PD-1 n=16, CpG (10 µg) + α-PD-1 n=20. **B)** Representative H&E stained histological images of mouse lungs from treatment groups: untreated LAM control, CpG (10 µg), α-PD-1, CpG (10 µg) + α-PD-1. Arrows indicate areas of necrosis. Scale bar = 500 µm. **C)** Quantification of LAM nodules and Inflamed tissue regions between treatment groups of mice at symptomatic endpoints from survival studies in (A). 3 sections analyzed per mouse. Statistical analysis was performed using log-rank test (survival studies) and one-way ANOVA (histological analysis). * p<0.05, ** p<0.01, *** p<0.001, **** p<0.0001.

CpG-ODN and anti-PD-1 therapy also resulted in reduced LAM nodule counts and lower abnormal tissue fractions, but did not reduce LAM nodule size as effectively as CpG-ODN and rapamycin combination therapy (**Fig. 8B-C**). In fact, combination therapy of CpG-ODN with checkpoint inhibition eventually resulted in larger average nodule sizes than those in untreated controls. This could be due to the class of CpG-ODN we selected for this study, as CpG-ODN-B has been shown to increase immune cell expression of PD-L1, and contribute to tumor escape and immune evasion (40). We observed signs of necrosis in the LAM nodules of checkpoint inhibitor and CpG-ODN combination therapy (**Fig 8C**) that were not present in nodules of untreated LAM controls or rapamycin and CpG-ODN combination therapy. Tumor necrosis is often associated with enhanced cytotoxic T-cell responses as a result of immune checkpoint inhibition.

To establish that combination rapamycin and anti-PD-1 therapy are compatible for treatment in LAM, we conducted a survival study and found the combination to be synergistic, increasing median survival to 77 days from 49 and 60 days for anti-PD1 and rapamycin respectively **(Fig. 9A).** Triple combination therapy of CpG-ODN with low-dose rapamycin and anti PD-1 therapy also proved to be synergistic, with a median survival of 95 Days **(Fig. 9B).** It is interesting to note that only the combinations that included both CpG-ODN and rapamycin resulted in the highest survival across all survival experiments (100 and 95 days), suggesting that the interaction between the two is the primary driver of these treatment outcomes.

**Figure 9.**
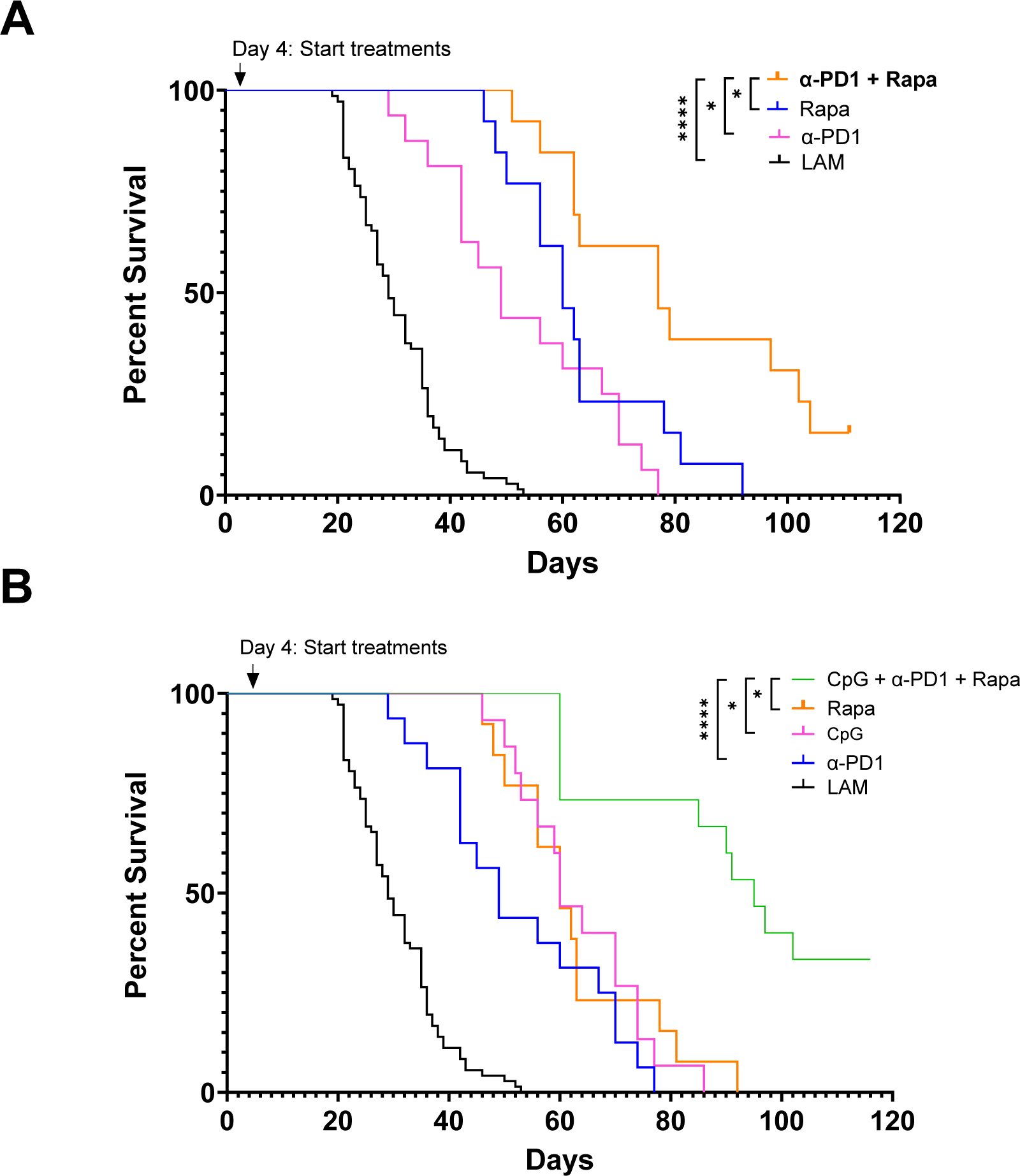
Triple Combination CpG-Rapamycin-ICI Therapy. **A)** Aggregate data from n>2 survival experiments LAM n=72, α-PD-1 n=16, Rapamycin n=15, Rapamycin + α-PD-1 n=16. **B)** Aggregate data from n>2 survival experiments LAM n=72, α-PD-1 n=16, CpG n=15, Rapamycin n=15, Rapamycin + CpG + α-PD-1 n=15. Statistical analysis of survival studies was performed using log-rank tests. * p<0.05, ** p<0.01, *** p<0.001, **** p<0.0001.

Taken together, our data suggests that that TLR9 activation may be beneficial as co-treatment with current clinical LAM treatment, rapamycin, and may also be combined with checkpoint inhibitors for increased efficacy.

## DISCUSSION

Our research offers preclinical evidence to support the benefit of adjuvant-mediated immunotherapy in the treatment of LAM. We found that intranasal treatment with CpG-ODN at early-stage disease increased survival in mice with LAM. This treatment slowed the progression of disease in the lungs and decreased the LAM nodule burden in treated mice. CpG-ODN treatment also increased the population of cytotoxic and effector CD8+ T cells and decreased the population of Treg cells in the LAM microenvironment. We found that pDC mediated induction of Th17 cells is in part responsible for CpG-ODN treatment efficacy. We also observed synergistic effects of both systemic checkpoint inhibition and low-dose rapamycin when combined with local CpG-ODN adjuvant therapy. This finding suggests that immune activating therapies, particularly combination therapies, could be an effective strategy to treat LAM.

Our finding that CpG-ODN treatment was not as effective for treating LAM at a more advanced stage of disease is consistent with evidence showing that patients diagnosed later in the course of their disease progression are not as responsive to therapy (41). It is possible that the stage of disease chosen for intervention in our mouse model was too advanced to be rescued by TLR activation or did not reflect the level of disease typical for a late stage human diagnosis. However, it is worth noting that even with early intervention CpG-ODN therapy, mice eventually develop lung lesions and respiratory symptoms. Additionally, it is unlikely that patients with advanced LAM would be able to tolerate the inflammatory effects of CpG-ODN in an already delicate tissue. CpG-ODN, like rapamycin, seems to be most effective when given early in the course of disease.

### pDCs and Th17s as Mediators of immunotherapeutic efficacy

Our data suggests that CpG-ODN efficacy is mediated in part by cytotoxic CD8+ T cells and Th17s that are activated through pDCs. CpG-ODN-stimulated pDCs have been shown to promote Th17 differentiation through increased secretion of cytokines like IL-6, IL-23 and TGF-β (42)), which can induce tumor regression in mice (43). In alignment with this, the depletion of pDCs from the lung microenvironment in our CpG-ODN-treated LAM mice results in decreased IL-6 and IL-23 production in the tissue and is correlated with decreased expansion of Th17 cells. Th17 cells are associated with both pro-tumor and anti-tumor immunity as a result of their ability to switch between pro-inflammatory Th17-Th1 and immunoregulatory Th17-Treg phenotypes. In anti-tumor immune responses, Th17 cells are involved in tumor infiltration (44) and the activation of cytotoxic CD8+ T cells(45). CpG-ODN stimulation has been shown to inhibit Tregs and promote Th17 differentiation in a mouse model of leukemia(46). Authors found that CpG-ODN tumor vaccine prolonged survival of mice with leukemia and suggest that this is due to both the reduction in Tregs and increased Th17 response. Additionally, Tregs can be skewed to generate IL-17 in the presence of pro-Th17 IL-6 and IL1β (47), and these Tregs are less effective at inhibiting effector Th17 and cytotoxic CD8+ T cells in autoimmune arthritis(48). Thus, our findings that Th17s are mediators of anti-LAM immunity after CpG-ODN treatment aligns with existing literature demonstrating the importance of inflammatory Th17s in anti-tumor immunity.

pDCs have also been implicated in mediating immune responses against viruses and cancer. pDCs have been shown to induce IFNy producing CD8+ and CD4+ T cells, crucial to resolving LCMV infection in the lungs(49). Additionally, in the lungs, pDCs were shown to affect Th17 differentiation in a mouse model of Bordetella pertussis infection. Interestingly, in B. pertussis infection pDCs reduced Th17 differentiation and blocking pDC produced IFNa resulted in reduced bacterial loads and increased Th17s(50). Overall, literature suggests that pDCs can have beneficial or detrimental effects, depending on the context. Our findings show that in LAM, depleting pDCs inhibits Th17 differentiation in helper T cells and reduces survival after CpG-ODN-mediated TLR9 activation, suggesting that pDCs play a beneficial role in this disease. We did not see increased levels of IL-17A cytokine secretion in response to CpG-ODN in LAM lungs (**Fig. E9)**. However, this could be due to the regulatory role that TLR9 activation appears to have on IL-17A production (51,52). More research is needed to understand the role of Th17 helper T cells in the immune response to CpG-ODN-treated LAM.

### Striking the balance between good/bad inflammation effects

A sophisticated balance of pro- and anti-inflammatory signals helps regulate immune responses in the body. Immunotherapies influence these pathways, tipping the scale towards inflammation to produce impressive anti-tumor responses. Our work here has shown that this balance is also disturbed in LAM and anti-LAM immunity can be activated using TLR9 adjuvant CpG-ODN. However, the inflammation induced can also lead to adverse effects and inflammatory toxicities. Choosing the right dose and treatment schedule to maintain equilibrium between treatment efficacy and runaway inflammation could be key to avoiding undesirable tissue damage and severe symptoms. Our finding that lower dose CpG-ODN therapy produces less inflammation in the lungs and increases survival in LAM mice despite the increase in nodule count (compared to higher dose CpG-ODN), suggests that too much inflammation may cause damage in an already delicate tissue environment like the LAM affected lungs and thus high levels of CpG-ODN have reduced treatment efficacy. A recent clinical trial looking at inhaled TLR9 agonist for the treatment of lung cancer reported several adverse events in patients all related to inflammatory immune responses (NCT03326752) (53), suggesting that TLR9 activation while potent may require more targeted delivery strategies that prevent exposure to all tissues. Our data suggest that this is true also for CpG-ODN treatment in LAM. A potential strategy to mitigate CpG-ODN side effects is to encapsulate CpG-ODN into a nanoparticle system that can deliver the TLR9 agonist directly to antigen presenting cells or the lymph nodes, where adaptive immunity is shaped. Encapsulation of CpG-ODN could reduce tissue-wide inflammation and thus reduce local toxicity effects, while maintaining the anti-LAM immune activation so crucial to inhibit LAM nodule proliferation. In addition, activation of stromal cells is correlated with tumor progression (54) and immunotherapy strategies inhibiting activated fibroblasts have been successful in promoting anti-tumor immunity in several preclinical cancer models (55–57). Because LAM nodules are composed of many cell types including fibroblasts that may contribute to LAM pathology (58–60), targeting activating treatments to immune cells in the lymph nodes while inhibiting activation of stromal cells in the tissue could help curb tissue remodeling (61).

### Potential for clinical translation of LAM immunotherapies

*Translating cancer immunotherapies to LAM:* Several immunotherapies are currently approved for treatment of lung cancer and may be considered for LAM. These include monoclonal antibodies against EGFR and VEGF/VEGFR pathways and immune checkpoint inhibitors targeting PD-1/PD-L1 or CTLA-4. Clinical trials for adoptive T cell therapy and lung cancer vaccines are ongoing (62) and several immunotherapies are approved for advanced lung cancers (63). Similarly, clinical trials for LAM therapies have explored the use of FDA-approved anti-angiogenic and anti-fibrotic therapies targeting pathways involved in the progression of lung cancer and idiopathic pulmonary fibrosis (Bevacizumab, Nintedanib, Saracatinib), but so far with limited success. Immune checkpoint inhibitors have not yet been tested in LAM patients, but there is promising research in mouse models to support this treatment strategy (17,18). Furthermore, our work showing TLR9 agonist treatment can enhance survival in murine LAM could be translated into the clinic in future, particularly since TLR9 agonists have been approved for several cancers and are explored in clinical trials for lung cancer. Given that our mouse model of LAM does not fully represent human LAM, we checked the presence of pDCs and TLR9 expressing antigen presenting cells in LAM tissues using the LAM Cell Atlas (64). The LAM Cell Atlas confirmed that TLR9 expressing pDCs are present in human LAM lung tissues, suggesting that a TLR9 stimulating treatment could be successfully translated to humans. Finally, results from lung cancer studies suggest that combination of immune checkpoint inhibitors and TLR9 agonists lead to successful anti-lung cancer immune re-activation. Our data suggests that this strategy could be successful in LAM as well and thus there is potential for designing LAM combination immunotherapies in the future.

*Combination therapy with existing LAM treatments:* Despite mTOR inhibition, we found that TLR-9 activation with CpG-ODN was synergistic with low-dose rapamycin therapy in our mouse model. This could be due to incomplete mTOR inhibition, as rapamycin reliably blocks mTORC1, but not mTORC2 (65). The effect of rapamycin on mTORC1 vs mTORC2 is dose dependent. Concentrations of rapamycin required to inhibit mTORC2 are much higher than those necessary to inhibit mTORC1 (66–68) and are not well tolerated for clinical intervention. Chronic inhibition of mTOR with rapamycin does eventually sequester enough free mTOR proteins to partially hinder a cell’s ability to assemble new mTORC2 complexes (69). However, low-dose rapamycin is not sufficient to completely suppress mTORC2 activity (70). In fact, evidence suggests that inhibition of mTORC1 can lead to activation of mTORC2 (71) and vice versa (72–74), potentially as a means to rescue proliferative functions through a parallel pathway.

In LAM patients, treatment with rapamycin is not curative but significantly slows the proliferation of LAM cells and disease progression (2,5,6). This is recapitulated in our model showing LAM nodules dispersed throughout the lungs at symptomatic endpoints despite increased survival when treating with low-dose rapamycin (**Fig. 21B**). Research points to activity of mTORC2 as a driver of persistent proliferation and nodule formation seen in LAM lungs despite inhibition of mTORC1 (75,76), and could explain incomplete patient responses to treatment with rapamycin alone.

Our finding that the benefit of CpG-ODN treatment was further enhanced with low-dose rapamycin is consistent with results showing that increased mTORC2 activity or mTORC1 inhibition work alongside TLR activation to generate pro-inflammatory immune responses. One research group found that mTORC2 activation increases TLR-4 induced proinflammatory cytokine production in human dendritic cells (82), and increases their capacity to prime T cells. Another study found that low-dose rapamycin alongside TLR activation increases TNF-α production in macrophages (83). In addition, mTORC1 inhibition significantly increases NF-KB activity and blocks the secretion of IL-10, an anti-inflammatory cytokine (77). (All TLRs, with the exception of TLR-3, signal through the MyD88-NFKB pathway to produce proinflammatory cytokines). This suggests that low-dose rapamycin treatment may not affect the ability of antigen presenting cells in the LAM microenvironment to respond to CpG-ODN and initiate inflammatory immune responses. In fact, low-dose rapamycin therapy has been shown to regulate and even increase the expansion of central memory T cells (78).

Rapamycin is the only FDA-approved treatment currently available to LAM patients, and its compatibility with CpG-ODN in our studies suggests this treatment strategy could merit consideration in clinical applications. It is possible that rapamycin’s cytostatic effect on LAM cell proliferation, combined with the immune infiltration mediated by CpG-ODN, alters the tissue microenvironment in the lungs in a way that synergistically restrains LAM nodule formation. More investigation into how this combination works will help in understanding how other immunotherapies may work in tandem with rapamycin. In addition, our finding that treatment with CpG-ODN alone is just as effective at increasing survival as low-dose rapamycin is further justification for its assessment in clinical trials. Given that 40% of LAM patients have partial or no response to treatment with rapamycin (79), TLR9 activation could offer these individuals a therapeutic strategy with similar benefits to standard of care.

### Conclusions

Our findings advance our understanding of potential immunotherapeutic strategies for LAM and their combined outcomes with standard of care. We present a novel approach (TLR9 activation through inhaled CpG-ODN treatment) that may be as effective as rapamycin and seems to have additive effects when combined. We have shown that TLR9 agonist CpG-ODN can enhance survival in a mouse model of LAM and is additive with anti-PD1 checkpoint inhibitor immunotherapy and low-dose rapamycin treatment.

We have also identified key players in the immune response generated by this response: plasmacytoid dendritic cells, and T cells, which could be important targets for future immunotherapy in LAM (**Figure 10**). This is important, because it shows that immune-modulation is possible in the LAM microenvironment, and works to significantly improve relevant outcomes in our preclinical models of disease. This research highlights a new potential area to develop for LAM treatments and combination immunotherapies, and has promising implications for the use of CpG-ODN as an adjuvant in the development of vaccines against LAM specific antigens when reliable markers for these disease cells can be identified.

**Figure 10.**
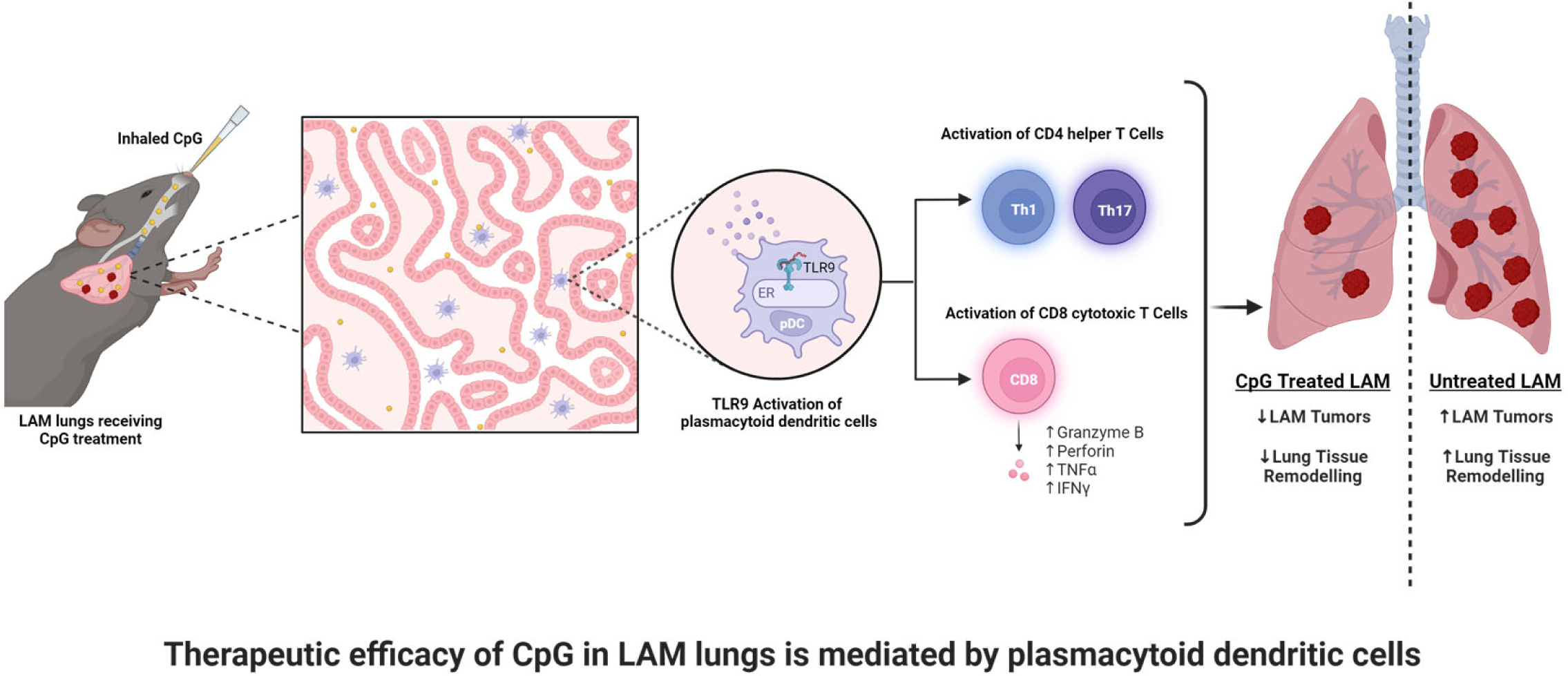
Summary of Findings.

