## Supplemental Figures for "Plasmacytoid dendritic cells mediate CpG-ODN induced increase in survival in a mouse model of lymphangioleiomyomatosis"

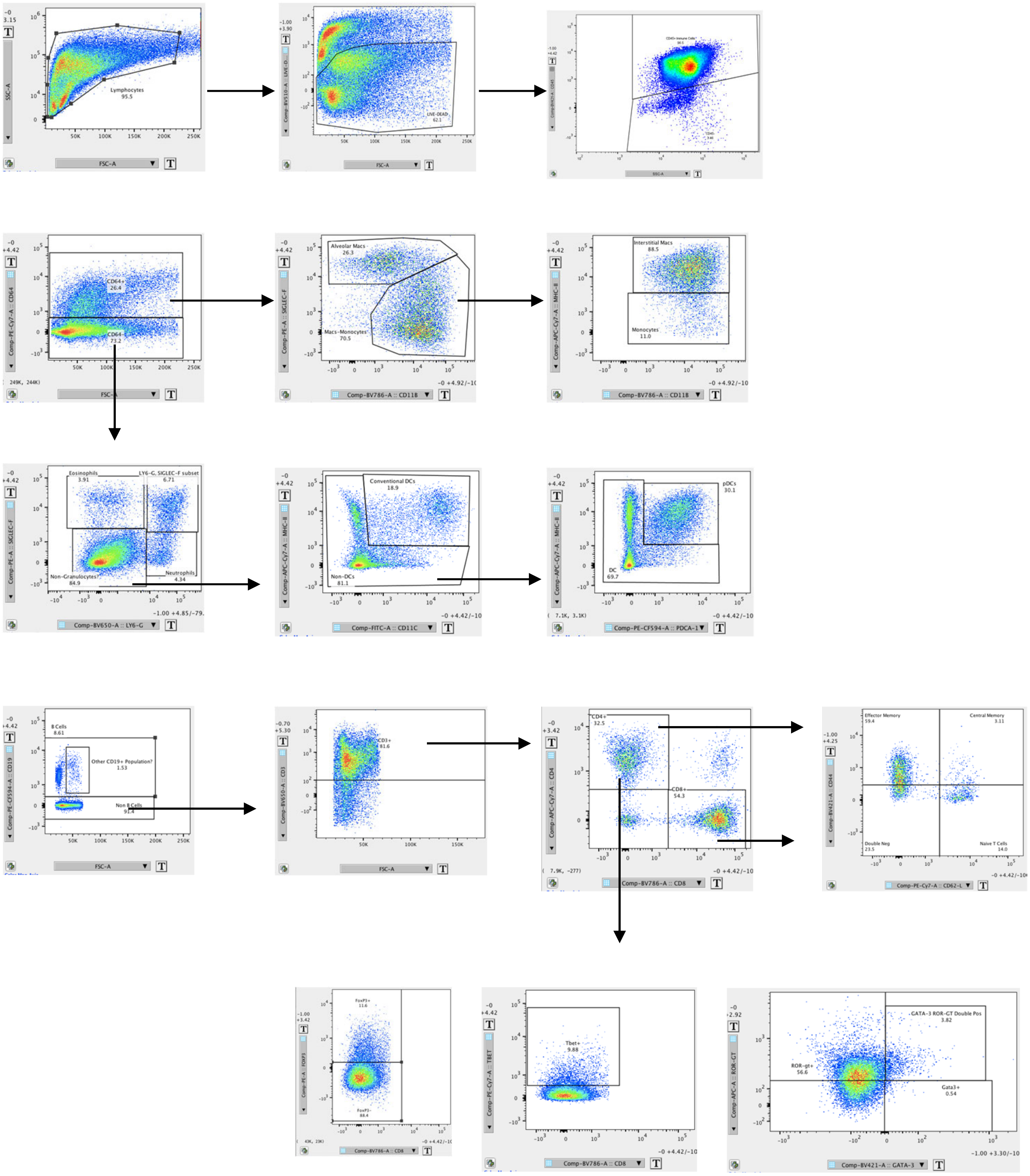


**Figure E1**: Flow Cytometry Gating Strategy

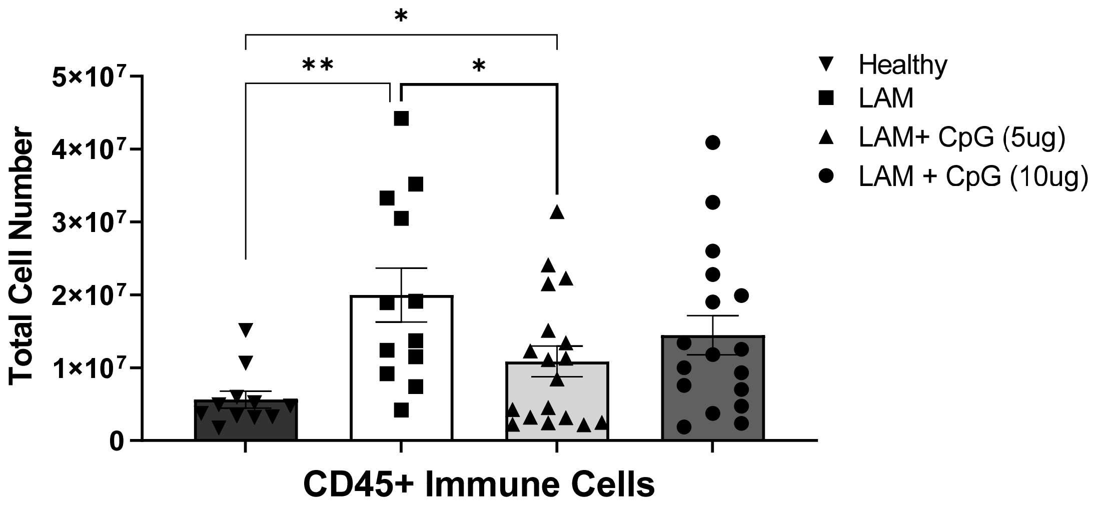


**Figure E2:** Total immune cell counts in lungs of mice on day 22 in early stage disease with or without CpG treatment. Data is representative of n=2+ experiments with n>4 mice. Statistical analysis was performed using two-way ANOVA. * $p<0.05$, ** $p<0.01$, *** $p<0.001$, **** $p<0.0001$.

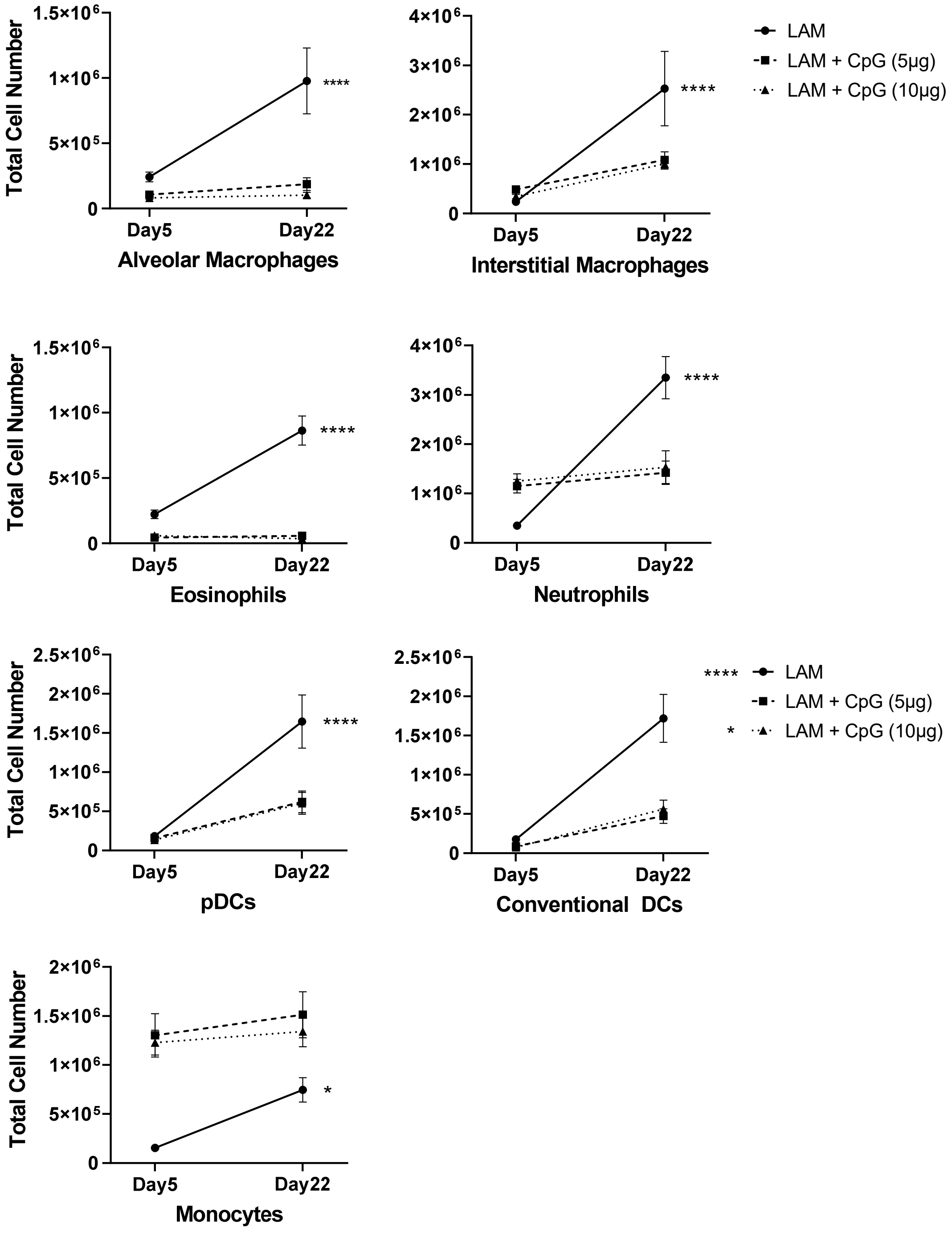


**Figure E3:** Total cell counts of granulocytes and antigen presenting cells on day 5 and 22 in early stage disease with or without CpG treatment . Data is representative of n=2+ experiments with n>4 mice. Statistical analysis was performed using two-way ANOVA. * $p<0.05$, ** $p<0.01$, *** $p<0.001$, **** $p<0.0001$.

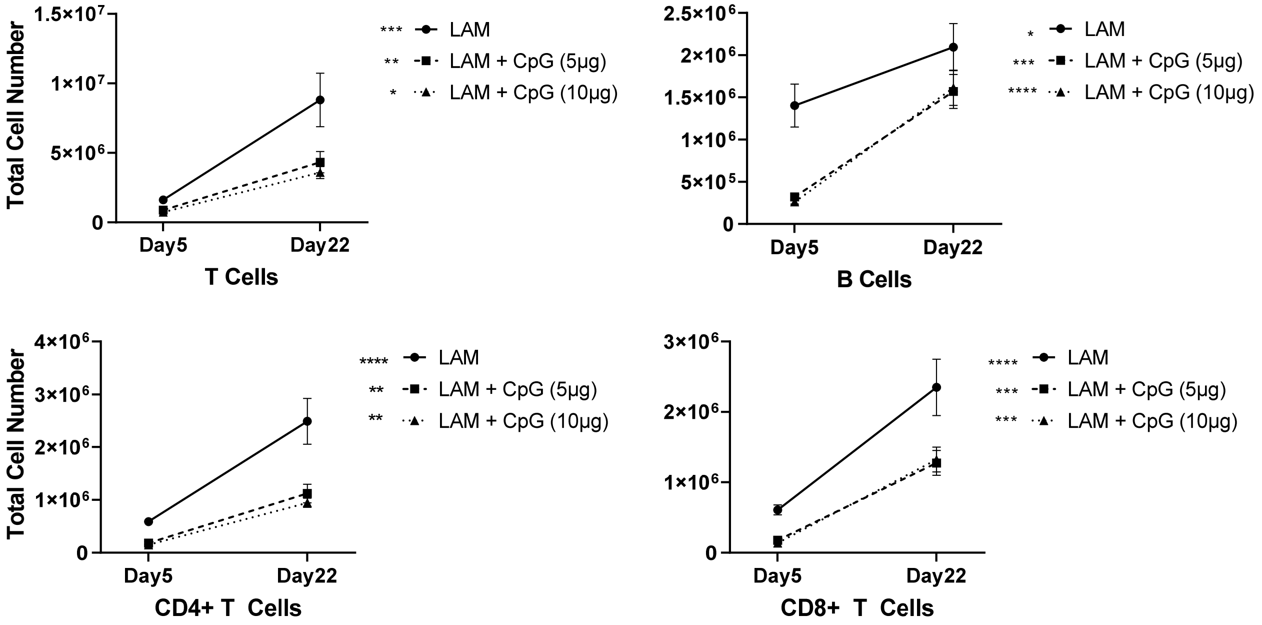


**Figure E4:** Total cell counts of T and B cells on day 5 and 22 in early stage disease with or without CpG treatment . Data is representative of n=2+ experiments with n>4 mice. Statistical analysis was performed using two-way ANOVA. * p<0.05, ** p<0.01, *** p<0.001, **** p<0.0001.

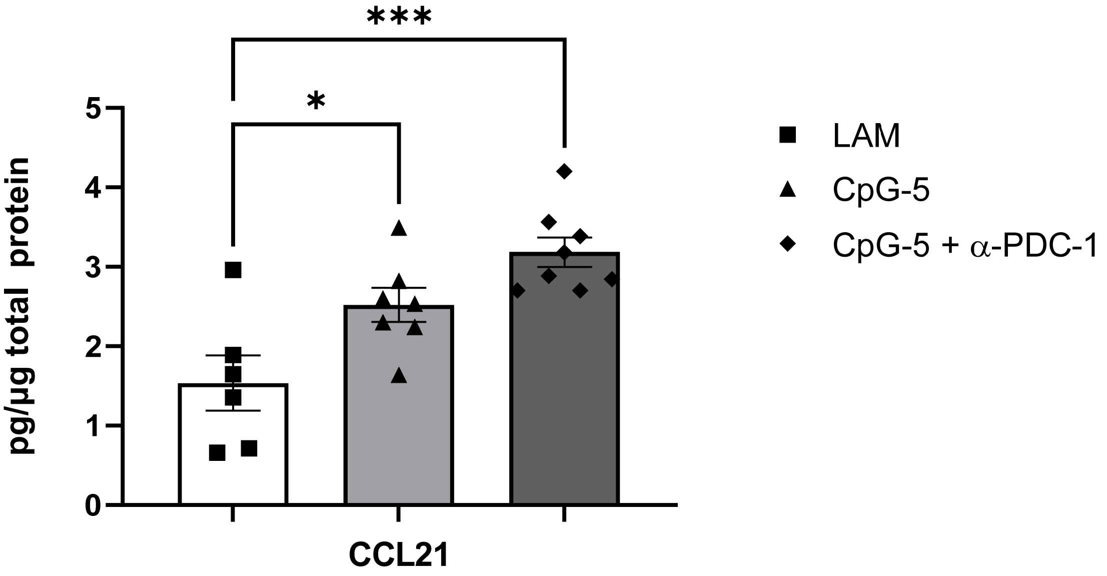


**Figure E5:** T cell chemoattractant, CCL21 in lung homogenate from CpG-treated mice with LAM and with pDC depletion. Data is representative of n=2+ experiments with n>4 mice. Statistical analysis was performed using ordinary one-way ANOVA. * $p<0.05$, ** $p<0.01$, *** $p<0.001$, **** $p<0.0001$.

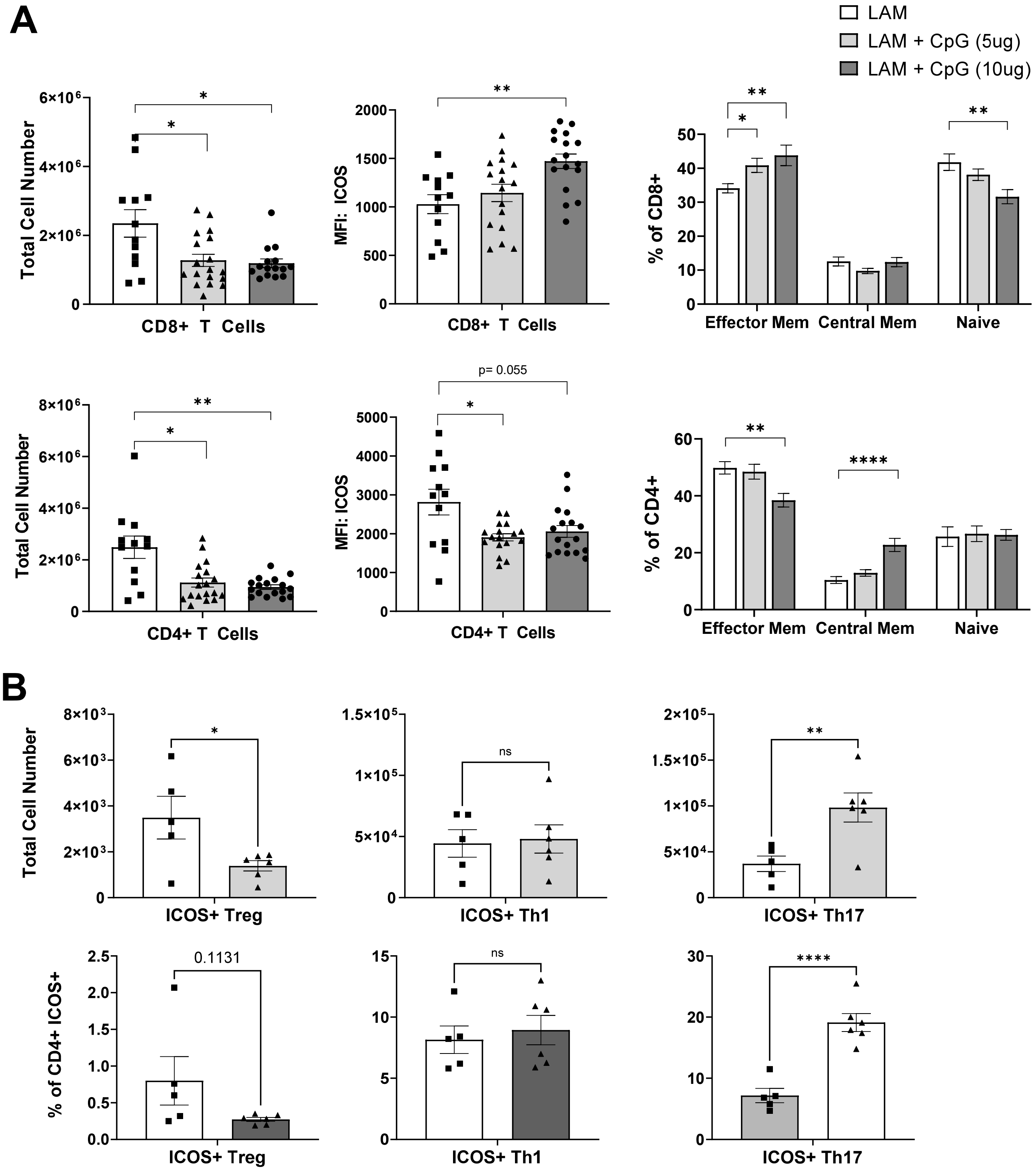


**Figure E6:** T cells after CpG treatment on day 22 in early stage disease with or without CpG treatment. **A)** Data shows CD4+ and CD8+ T cell numbers, ICOS expression, and cell population distribution between effector memory (CD44+CD62L-), central memory (CD44+CD62L+) and naïve subsets (CD44-CD62L+). **B)** Data shows ICOS+ regulatory, Th1, and Th17 CD4+ T cells: numbers and %CD4+. Data is representative of n= 2-3 experiments and n>10 mice. Statistical analysis was performed using one-way ANOVA (A) or student's t test (B). * $p<0.05$, ** $p<0.01$, *** $p<0.001$, **** $p<0.0001$.

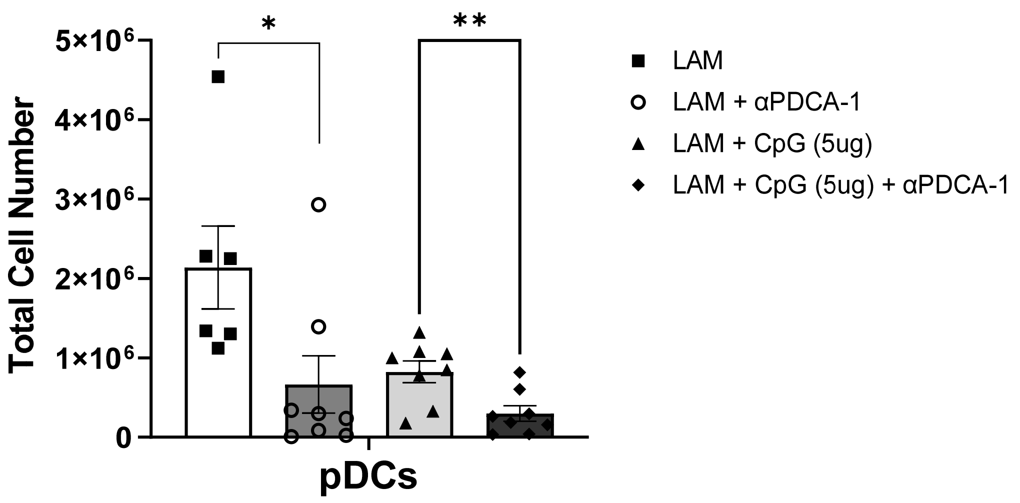


**Figure E7:** Total pDC counts on day 22 in early stage disease with or without CpG treatment. Data is representative of n=2+ experiments with n>4 mice. Statistical analysis was performed using two-way ANOVA. * $p<0.05$, ** $p<0.01$, *** $p<0.001$, **** $p<0.0001$.

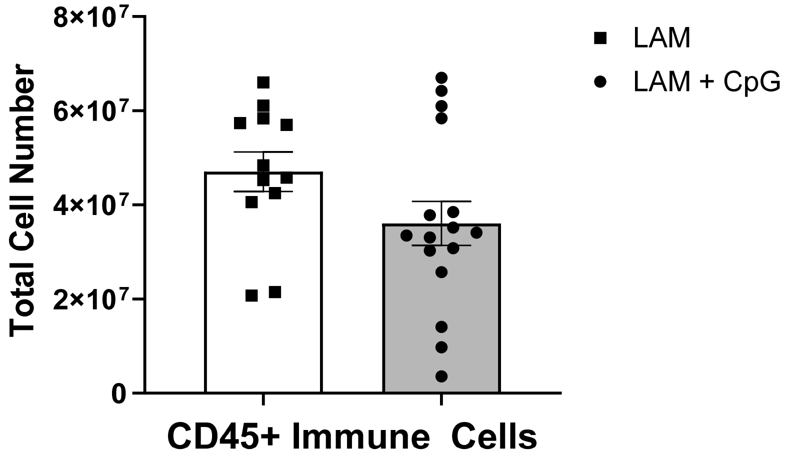


**Figure E8:** Total immune cell counts on day 15 after late intervention CpG treatment on day 14. Data is representative of n=2+ experiments with n>4 mice. Statistical analysis was performed using two-way ANOVA. * $p<0.05$, ** $p<0.01$, *** $p<0.001$, **** $p<0.0001$.

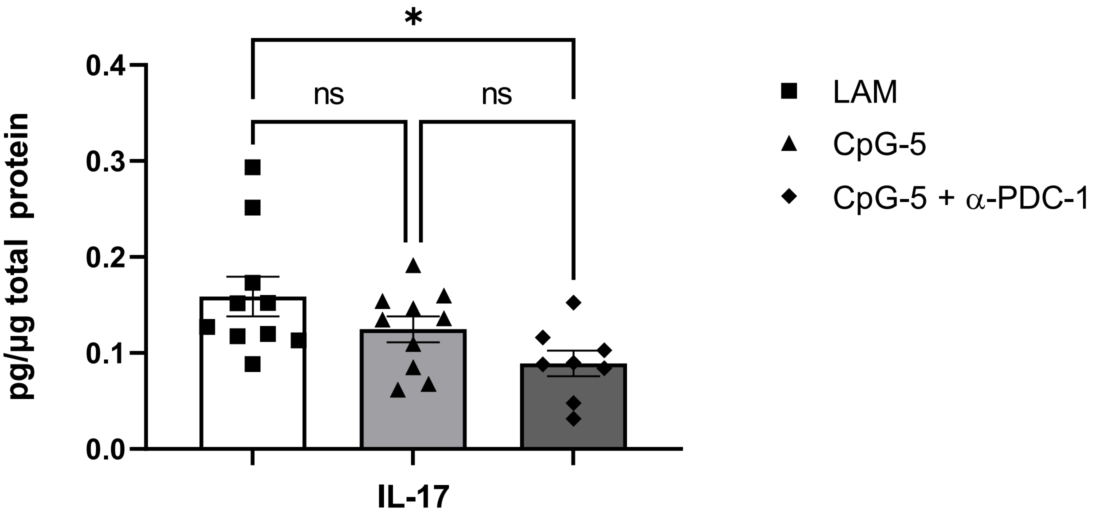


**Figure E9:** IL17 A/F in lung homogenate from CpG-treated mice with LAM. Data is compiled from n=2 experiments with n>4 mice. CpG-treated mice with LAM. Statistical analysis was performed using ordinary one-way ANOVA. * p<0.05, ** p<0.01, *** p<0.001, **** p<0.0001.
